# Emergence of bluetongue virus serotype 3 in western Germany, October 2023, and *ad-hoc* monitoring in *Culicoides* biting midges

**DOI:** 10.1101/2024.02.26.582175

**Authors:** Anja Voigt, Helge Kampen, Elisa Heuser, Sophie Zeiske, Bernd Hoffmann, Dirk Höper, Mark Holsteg, Franziska Sick, Sophia Ziegler, Kerstin Wernike, Martin Beer, Doreen Werner

**Author notes:** Corresponding author: Martin Beer, Friedrich-Loeffler-Institut, Federal Research Institute for Animal Health, Suedufer 10, 17493 Greifswald – Insel Riems, Germany.

## Abstract

Bluetongue virus serotype 3 emerged in October 2023 in Germany, where Schmallenberg virus is enzootic. BTV-3 was detected in a pool of *Culicoides* biting midges at the same time as infections were reported from ruminants. Schmallenberg virus was frequently found in vector pools, reflecting the epidemiological situation.

**One sentence summary line:** Coinciding with the emergence and low-level circulation of bluetongue virus serotype 3 in ruminants in Germany in October 2023, the virus was detected in a pool of *Culicoides* biting midges of the Obsoletus Group, while enzootic Schmallenberg virus was found in numerous midge pools.

## Main text

Bluetongue virus (BTV), an orbivirus of the *Sedoreoviridae* family, may cause epizootic disease in domesticated and wild ruminants (1). Bluetongue (BT) is a WOAH listed disease and regulated within the European Union (EU) in accordance with Regulation (EU) 2016/429 (“Animal Health Law”) and its delegated regulations (2), with outbreaks requiring trade restrictions in EU member states to prevent the etiologic agent from spreading. BTV-3 emerged for the first time in continental Europe in early September 2023, when clinical disease was observed on four sheep farms in the Netherlands. By mid-October, more than 1,000 outbreaks had been detected throughout the Netherlands, increasing to 5,884 by mid-December (3). Meanwhile, BTV-3 has also reached Belgium and the UK (4, 5). In contrast to the newly emerging BTV-3, the orthobunyavirus Schmallenberg virus (SBV) is enzootic in continental Europe since its first appearance near the German-Dutch border in 2011 (6). BTV and SBV share major epidemiological characteristics, as both are transmitted by *Culicoides* biting midges (Diptera: Ceratopogonidae) and affect mainly ruminants (1, 7).

### The Study

Shortly after Germany was officially declared free of BTV (BTV-8) in June 2023 (7), the first German case of BTV-3 was confirmed on 12 October in a sheep in the German district of Kleve, close to the Dutch border. Until February 15^th^, 2024, 38 additional BTV-3 cases were reported from sheep and cattle farms in the German federal states of North Rhine-Westphalia and Lower Saxony (Fig. 1). An isolate was obtained from a BTV-3 positive blood sample and further characterized on both *Culicoides* cells (KC cells) and baby hamster kidney cells (BHK cells). Whole genome sequencing resulted in a nearly complete genome sequence (available from the INSDC databases under project ID PRJEB72862). Overall, the obtained genome is 99.94% at the nucleotide level and 99.95% at the amino acid level, respectively, identical with the sequence of a recent BTV-3 isolate from the Netherlands (OR603992.1, OR603993.1, OR603994.1, OR603995.1, OR603996.1, OR603997.1, OR603998.1, OR604000.1, OR603999.1, OR604001.1).

**Figure 1:**
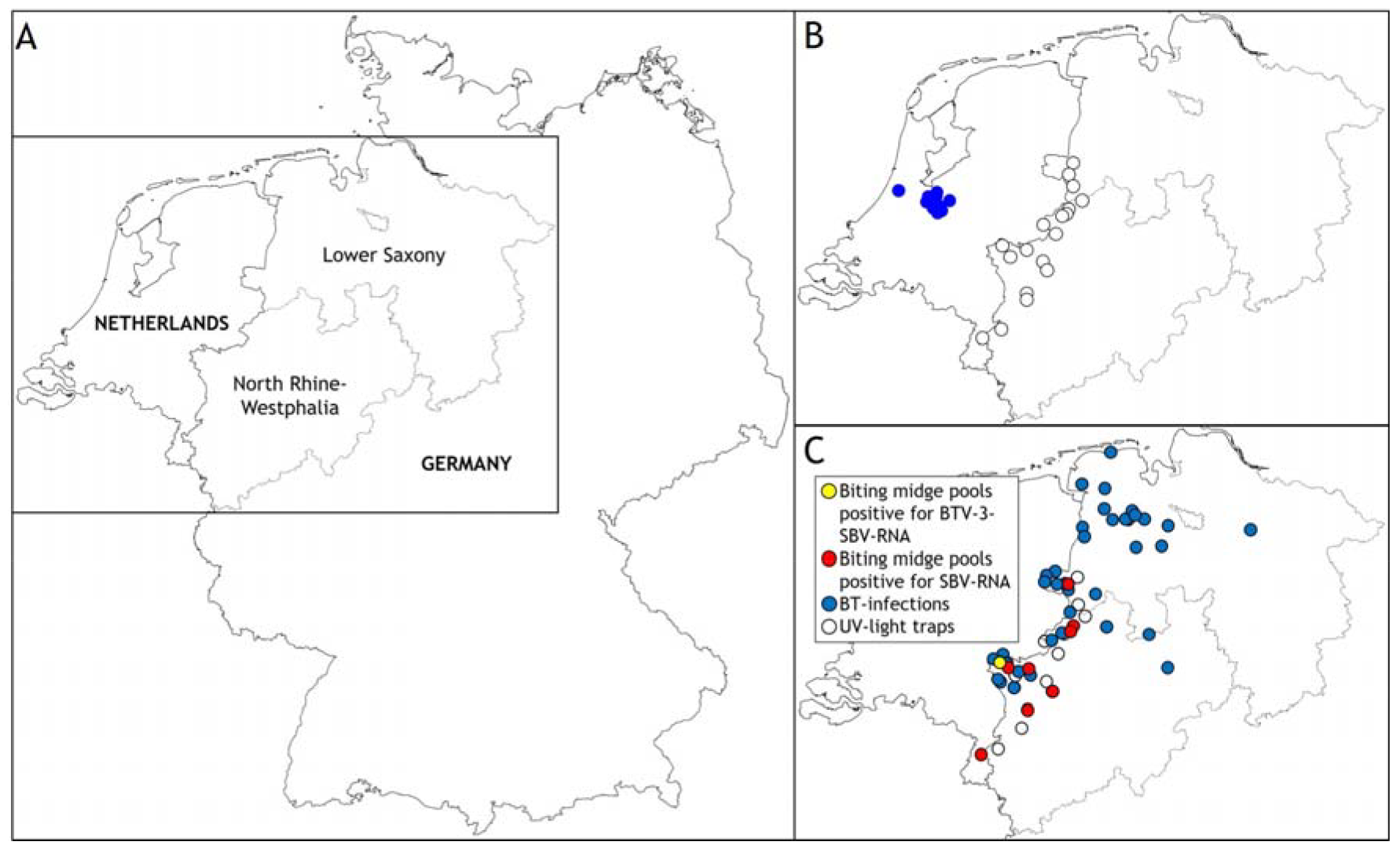
(A) Overview Germany – Netherlands – German federal states of North Rhine Westphalia and Lower Saxony. (B) Dutch BTV-3 cases (blue dots) as of 08 Sept 2023 and locations of UV-light-traps along the German-Dutch border (white dots). (C) BT infections in ruminants (blue dots) as reported to the German animal disease reporting system as of 15 Feb 2024 and geographic assignment of BTV-3-/SBV-RNA- (yellow dot) and SBV-RNA- (red dots) positive biting midge pools.

Following the emergence of BTV-3 in the Netherlands, i.e. before any clinical suspicions had been announced in ruminants in Germany, biting midge traps were installed in animal stables in western Germany to collect putative BTV-3 vectors and test them for virus infection. Between September 24 and 26, UV-light-traps were set up in cattle, sheep and goat sheds on 18 German farms along the Dutch border (North-Rhine Westphalia and Lower Saxony). The traps were placed close to the animals and were protected from wind and rain. Insects were sampled daily until 9 or 11 November, depending on the location, and then once a week for 24 hours. Collected biting midges were morphologically identified according to ‘Obsoletus Group’, ‘Pulicaris Complex’ and ‘other *Culicoides’*. Obsoletus Group and Pulicaris Complex, which are considered to contain the main virus vectors in Europe (8), were screened in pools of up to 50 individuals for RNA of BTV and epizootic haemorrhagic disease virus (EHDV), another culicoid-borne orbivirus that recently emerged in Europe (9). To include a virus that is enzootic in the ruminant population in the region, and that is consequently expected to be present in the insect vectors at detectable prevalences, biting midges were also tested for Schmallenberg virus (SBV). The SBV-specific RT-qPCR system used has been described earlier (10), and for BTV and EHDV detection a multiplex RT-qPCR was used (11). Pools that tested positive for BTV were subsequently analyzed by a BTV-3-specific RT-qPCR (12). Biting midge pools positive for viral RNA were retrospectively examined for the biting midge species contained in the pools (13, https://biorxiv.org/cgi/content/short/2024.01.23.576915v1).

In a first approach, 1,603 biting midge pools collected at nine sites from 26 September to 09 November were tested for BTV, SBV and EHDV, with the number of pools per site ranging from 27 to 466. One pool collected October 12 in the district of Kleve (Fig. 1) was tested positive for BTV-RNA (quantification cycle [Cq] value: 35.6), and was subsequently confirmed as BTV-3 positive (Cq 37.5). The pool consisted of a mixture of *C. obsoletus* clade O1, *C. scoticus* and *C. chiopterus*. Another pool with Obsoletus Group biting midges collected on the same day and at the same site tested positive for SBV. In addition, SBV was detected in 534 further samples of this 6.5-week-period from all nine locations, one site in Lower Saxony in the district of Grafschaft-Bentheim and eight sites in North Rhine-Westphalia (three in the district of Kleve, two in the district of Wesel, two in the district of Borken and one in the district of Heinsberg) (Fig. 1; Appendix). With the exception of two Pulicaris Complex pools, all SBV-RNA-positive pools belonged to the Obsoletus Group (for taxon specification see Appendix). No pool was found positive for both BTV- and SBV-RNA, and all pools tested were negative for EHDV.

## Conclusions

BTV-3 was first confirmed in Germany on 12 October 2023 in a sick sheep, and further spread was detected in two German federal states until winter 2023/2024. The isolated BTV-3 is almost identical to the virus of the outbreaks in the Netherlands. A pool of biting midges of the Obsoletus Group – collected on the day of the first BTV-3 confirmation in Germany – was found positive for BTV-3-RNA on a cattle farm in the same district. The detection of the virus in their putative vectors confirms an ongoing transmission cycle at that time, albeit circulating at a very low level, as only a single insect pool tested positive and only single animals were positive in infected farms. In contrast, the SBV genome was found in numerous *Culicoides* pools, reflecting the situation in ruminant populations, as SBV-infected cattle, sheep and goats have been reported in Germany since 2011, albeit with varying prevalence between years (6). The Cq values of the SBV RT-qPCR in some of the investigated *Culicoides* pools (Appendix) indicate substantial virus loads, reflecting extensive regional SBV circulation in autumn 2023. While SBV has become enzootic in Germany, the new outbreak of BTV-3 is expected to continue, intensify and spread with the seasonal onset of biting midge activity in spring 2024.

## Acknowledgments

We would like to thank all farmers who supported the study and volunteered in attending biting midge traps. The study was funded by the German Federal Ministry of Food and Agriculture (BMEL) through the Federal Office for Agriculture and Food (BLE), grant numbers 28N207601 and 28N207602, as well as EU Horizon 2020 program project Versatile Emerging infectious disease Observatory (VEO), grant number 874735.

## Author biography

Anja Voigt is a Ph.D. student at the Leibniz-Centre for Agricultural Landscape Research, Muencheberg, Germany. Her research interests include behavior and habitat binding of biting midge vectors.

